# Characterization of Vif domains that mediate Feline Immunodeficiency Virus antagonism of APOBEC3-H and APOBEC3-CH restriction

**DOI:** 10.1101/144113

**Authors:** Olivia L. Sims, Ernest L. Maynard, Eric M. Poeschla

## Abstract

Feline immunodeficiency virus (FIV) Vif mediates degradation of two anti-lentiviral feline APOBEC3 (fA3) proteins, fA3Z3 and fA3Z2bZ3. HIV-1 Vif targets the restriction factor human APOBEC3G (A3G, hA3Z2g-Z1c) for proteasome degradation to mediate viral evasion. Despite this similarity, FIV and HIV-1 Vif share limited homology. Vif binds hA3Z2g-Z1c through its N-terminal region, while its C-terminal region binds to an E3-ligase complex containing Cullin5 and Elongin B/C. Further, HIV-1 Vif contains critical domains in its C-terminus, including an adjacent BC box, the only shared domain between FIV and HIV-1 Vif, and a non-classical zinc finger (HCCH) domain. Felid lentivirus Vif, however, contains a highly conserved KCCC motif. While both Vifs have evolved to counteract select A3 antiretroviral proteins, the FIV Vif domains necessary to target fA3s for degradation are incompletely understood. To identify these domains, we used the well-characterized HIV-1 Vif domains to show that distinct mutations within the BC box of FIV Vif prevent fA3Z3 and fA3Z2bZ3 degradation and reduce virion infectivity. We also found that mutating any single residue in the KCCC motif blocked fA3 targeting and impaired FIV infectivity and replication. These mutations also failed to disrupt the FIV Vif and Cullin5 interaction. Further, we showed that, in contrast to the HCCH domain in HIV-1 Vif, the KCCC domain of FIV Vif does not bind zinc. However, unlike HIV-1 Vif, FIV Vif (C36 isolate) reduces intracellular levels of co-expressed Cullin5 proteins, a novel finding. Our results reveal important C-terminal residues in FIV Vif and show that the BC box and KCCC regions are critical for fA3 degradation, infectivity, and spreading replication.

## Introduction

Lentiviruses cause chronic degenerative diseases in primate, ungulate, and feline species. The current AIDS pandemics in *Homo sapiens* and *Felis catus* are caused by human immunodeficiency virus type-1 (HIV-1) and feline immunodeficiency virus (FIV), respectively. All primate and non-primate lentiviruses encode a Vif protein, except the equine virus (1), that is essential for viral replication. HIV-1 Vif counters the antiviral activity of human APOBEC3G (A3G, hA3Z2g-Z1c) (2–11). In the absence of Vif, hA3Z2g-Z1c incorporates into virion particles of the virus-producing cell, where it inhibits viral replication by cytidine deamination of the viral minus–strand cDNA. This mechanism produces a plus-strand G-to-A hypermutation and interference during reverse transcription (4, 12–15).

A3 proteins, such as hA3Z2g-Z1c, are targeted to the proteasome for degradation (2, 16–22) via a Vif-dependent mechanism. The N-terminus of HIV-1 Vif, the ^14^DRMR^17^ and ^40^YHHY^44^ motifs (23–26), mediates the interaction with hA3Z2g-Z1c. However, the C-terminal region of Vif coordinates recruitment of a cellular E3 ligase complex consisting of Cullin5 (Cul5), Elongin C, Elongin B, an E2 ligase, and a Ring box (Rbx) protein (2, 5, 6, 18, 27). Recruitment of the E3 ligase occurs via several evolutionarily conserved domains in Vif, including a non-classical zinc-finger domain that binds to Cul5, SOCS (suppressors-of-cytokine-signaling-like domain) box, BC box, that interacts with Elongin C (2, 5–9, 15, 28–30), and a PPLP domain that interacts with Elongin B (1, 6, 31). While there are seven different human Cullin proteins (32, 33), HIV-1 Vif selectively interacts with Cul5 to stabilize the E3 ligase complex (2).

The half-life of both HIV-1 and simian immunodeficiency virus (SIV) Vif is stabilized and extended by core-binding factor-β (CBF-β) (34), which promotes the interaction between HIV-1/SIV Vif and Cul5 needed to form the E3-ligase complex (27, 35–39). The interface that mediates CBF-β binding to and stabilization of Vif were solved using the Vif/CBF-β/Cul5/Rbx2/EloBC and, separately, the Vif/CBF-β/N-terminal hA3Z2g-Z1c/EloBC complexes with X-ray crystallography and NMR spectroscopy, respectively (37, 40). CBF-β forms multiple interactions with the HIV-1 Vif N- and C-terminus, particularly with the BC box and HCCH domains (40–42). Mutating HCCH residues prevents the HIV-1 Vif/CBF-β/hCul5 interaction and significantly suppresses hA3Z2g-Z1c antiviral activity (41).

Non-primate mammals also encode antiviral APOBEC3 proteins (11, 43–50). Two *Felis catus* A3 proteins, fA3Z3 (fA3H) and fA3Z2bZ3 (fA3CH), restrict Δvif FIV (11, 43–46), a FIV vif-deletion mutant. FIV Vif promotes HIV-1 replication by mediating degradation of fA3Z3 or fA3Z2bZ3 when expressed in *cis* (45, 46) or *trans* (45) in Crandall-Rees Feline Kidney (CrFK) cells. FIV Vif also recruits an E3-ligase complex containing Cul5, EloC, and EloB (51). Although FIV Vif lacks the HCCH motif, a region adjacent to the BC box contains a strongly conserved KCCC motif (Fig. 1A) (45) that (45, 51) may serve as an atypical putative zinc-binding domain. Other lentiviral proteins contain an atypical zinc finger, such as the HIV-1 nucleocapsid (NC), that is important for transferring DNA intermediate strands during reverse transcription (52–54). Additionally, FIV Vif interacts with human and feline Cul5 (fCul5) (51); however, the residues governing the interaction between Vif and Cul5 are different between HIV-1 and FIV Vif. Indeed, replacing the two most C-terminal cysteine residues in the KCCC motif of FIV Vif with serine (KCSS) no longer rescues ΔVif HIV-1 infectivity in the presence of fA3 proteins, despite maintaining the interaction between FIV Vif and Cul5 (51).

**Fig 1.**
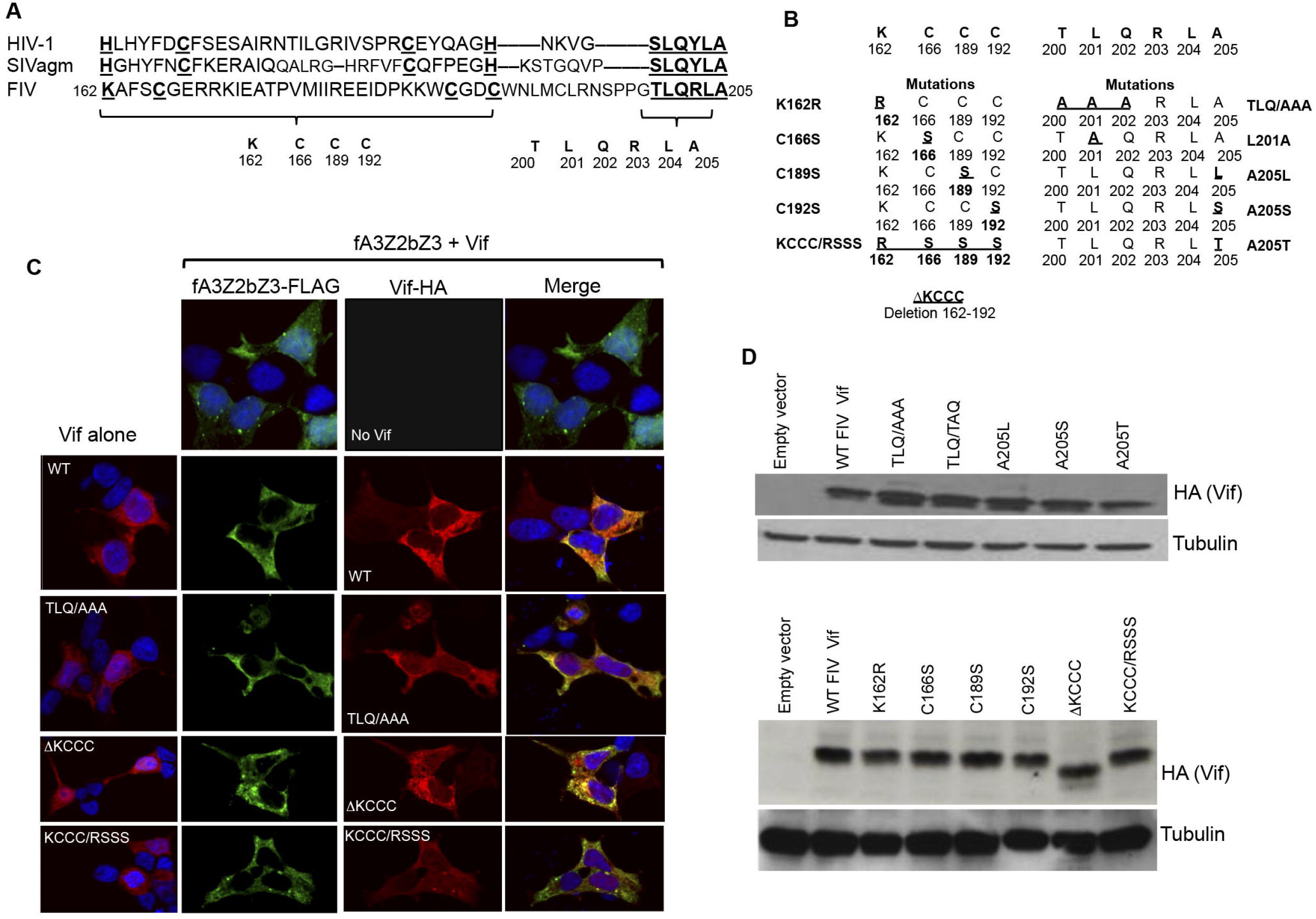
Vif protein alignments, mutations, cellular locations and expression levels. *A*, Amino-acid alignment of the C-terminus of HIV, SIVagm, and FIV Vif. *B*, FIV Vif C36 mutants. *C*, 293T cells were transfected with HA-tagged wild-type (WT) or mutant Vif with or without FLAG-tagged fA3Z2bZ3. Cells were subjected to immunofluorescence with antibodies to the respective epitope tags to visualize the cellular distribution of FIV Vif or fA3Z2bZ3 proteins alone (top row, left column) or together. Cellular DNA was stained with DAPI. *D*, Immunoblots demonstrating equivalent expression of HA-tagged FIV Vif mutants.

The zinc chelator PTEN (N, N, N’, N’-tetrakis-(2-pyridylmethyl)-ethylenediamine) further distinguishes FIV from HIV-1. PTEN disrupts the interaction between HIV-1 Vif and hCul5, but not between FIV Vif and fCul5, suggesting that FIV Vif might not be a bona fide zinc-binding protein (9, 29, 51). However, PTEN can chelate iron and copper (albeit it has a greater affinity for copper) and alter their cellular levels (55–57). While earlier data suggested that FIV Vif does not bind zinc, biochemical data using purified proteins were needed to verify this notion. Because HIV-1 NC has a non-canonical zinc motif, we posited that the KCCC domain in FIV Vif might be the major zinc-binding site and may mediate a functionally important Vif-Cul5 interaction. To test this hypothesis, we interrogated the biomolecular interactions between FIV Vif and Cul5 using isothermal titration calorimetry with and without zinc. Our studies were designed to decipher whether FIV Vif is a zinc-binding protein and whether additional determinants in the BC box and KCCC domain are essential for this interaction.

## Materials and Methods

### Cells

293T and CrFK cells were obtained from the American Type Culture Collection (ATCC). Cells were cultured in high-glucose Dulbecco’s Modified Eagle Medium (DMEM) with 10% fetal bovine serum, 1% penicillin/streptomycin, and L-glutamine.

### Plasmids

The expression plasmid containing codon-optimized wild-type (WT) FIV Vif C36 tagged with a hemagglutinin (HA) epitope (FIV C36 Vif-HA) was generated as previously described (45). Primer pairs (Table 1) were used for site-directed mutagenesis within FIV C36 HA-tagged Vif using sense and antisense primers to generate the following FIV Vif mutants: TLQ/AAA, TLQ/TAQ, A205L, A205S, A205T, K157R, C161S, and C187S. To generate the FIV Vif KCCC/RSSS mutant, a multisite-directed mutagenesis kit (Invitrogen) was used with two sense primers outlined in the primer table. Overlap-extension PCR deleted the KCCC domain using FIV C36 Vif-HA as a template with the outer sense (OS) and inner antisense (IAS) primers to generate the first PCR product. The second PCR reaction used the inner sense (IS) and outer antisense (OAS) primers. Combined PCR products were amplified with OS and OAS primers to generate ΔKCCC FIV C36 Vif-HA. PCR products were ligated into p1012 using EcoRI and KpnI restriction sites. The expression plasmid containing VR1012 human Cul5 Myc was a generous gift from Dr. Xiao Fang Yu at John Hopkins University School of Public Health (2).

**Table 1.**
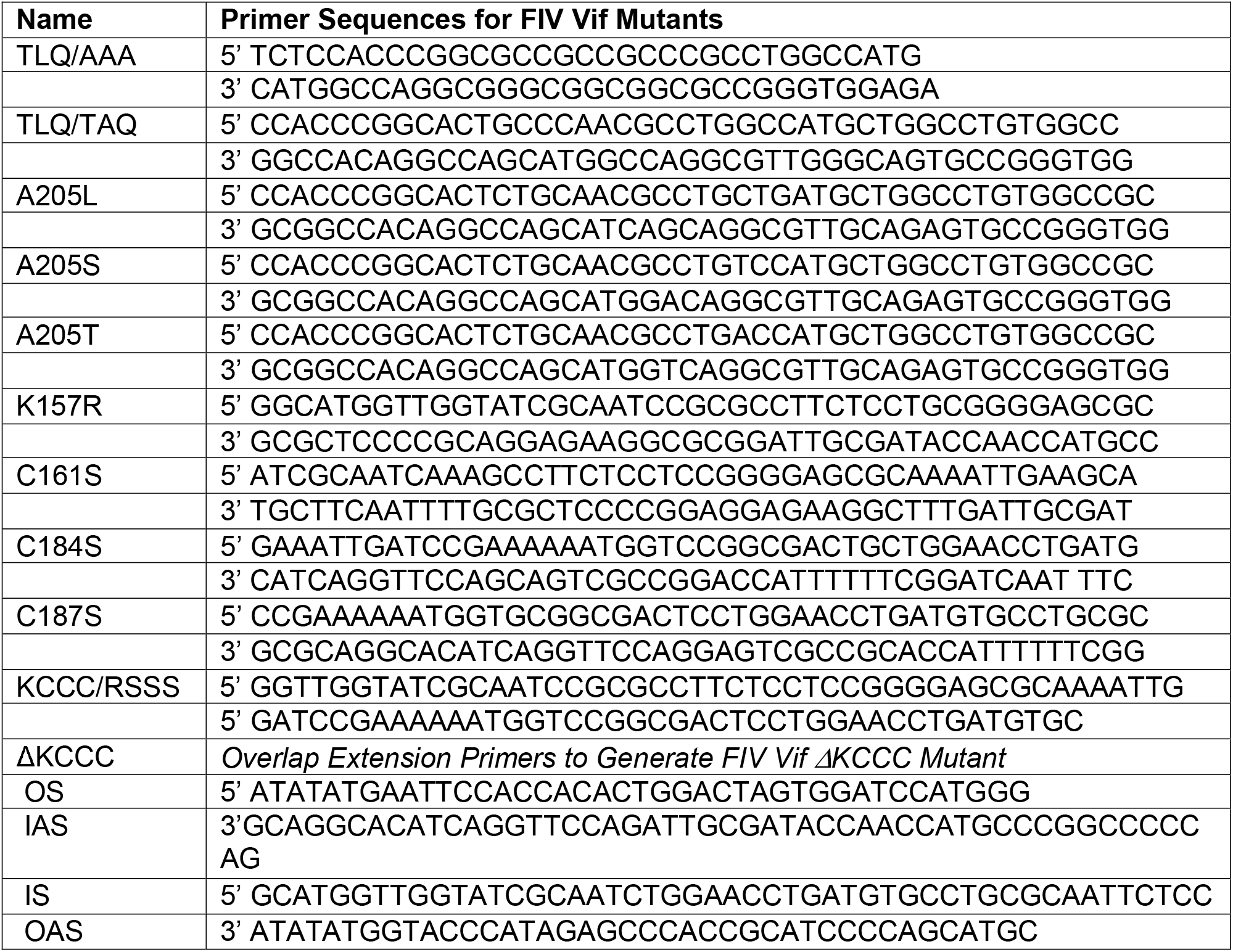
Primer Sequences to Generate FIV Vif Expression Plasmids.

Construction of expression plasmids containing fA3Z3 and fA3Z2bZ3 FLAG was previously described (45). The VR1012 feline Cul5 Myc was a generous gift of Dr. Yu (51). All expression plasmids were verified by Sanger DNA sequencing.

### Construction of Vif-Mutant Infectious Molecular Clones

#### Generation of 34TF10 C184S_187S Vif

The human cytomegalovirus driven infectious molecular clone 34TF10 (CT5) (58) containing a functional Orf2 was subcloned into pcDNA3.1(-) using EcoRI and KpnI (pcDNA3.1-CT5). pcDNA3.1-CT5 has a partial fragment of Gag and full length Pol, Vif, and Orf2 genes (hereafter referred to as pcDNA3.1-CT5Vif) (58). To introduce the C184_187S mutation into pcDNA3.1(-) CT5Vif, the OS and IAS primers were used to generate the first PCR amplicon. The second PCR product was synthesized with OAS and IS primers. Combined PCR products were amplified with the OS and OAS primers, and the overlap product was inserted into pCT5 with AccIII and KpnI.

#### Creation of KCCC/RSSS

To construct KCCC/RSSS, the C184_187S amplicon described above was ligated into pcDNA3.1(-) empty vector using EcoRI and KpnI. To generate KCCC/RSSS Vif, the pcDNA3.1-C184S_187S Vif plasmid and OS and IAS primers were used to amplify the first PCR product. The pcDNA3.1-C184_187S plasmid and OAS primer and the IS primers generated the second PCR amplicon. Combined PCR products were amplified using OS and OAS primers to generate the CT5 Vif KCCC/RSSS. The KCCC/RSSS PCR product was ligated into CT5 with AccIII and KpnI.

#### Creation of 34TF10 TLQ/AAA Vif

To generate the TLQ/AAA Vif mutant, the pcDNA 3.1-CT5 Vif plasmid and the OS and IAS primers were used to amplify the first PCR product. The pcDNA3.1-CT5 Vif template and OAS and IS primers were used to amplify the second PCR product. Combined PCR products were amplified with OS and OAS primers to generate the TLQ/AAA amplicon. The TLQ/AAA PCR product was ligated into CT5 with AccIII and KpnI.

#### Creation of 34TF10 ΔKCCC_TLQ/AAA Vif

To generate the ΔKCCC_TLQ/AAA Vif mutant, the CT5 TLQ/AAA amplicon was ligated into pcDNA3.1(-) using EcoRI and KpnI. The CT5 TLQ/AAA CT5 Vif template and OS and IAS primers were used to amplify the first PCR product. The second PCR product was amplified using the CT5 TLQ/AAA CT5Vif template DNA with OAS and IS primers. Combined PCR products were amplified using OS and OAS primers to generate the full-length ΔKCCC_TLQ/AAA Vif amplicon. The primers used to generate mutants with overlap-extension PCR are listed in Table 2. All mutants were verified by Sanger DNA sequencing.

**Table 2.**
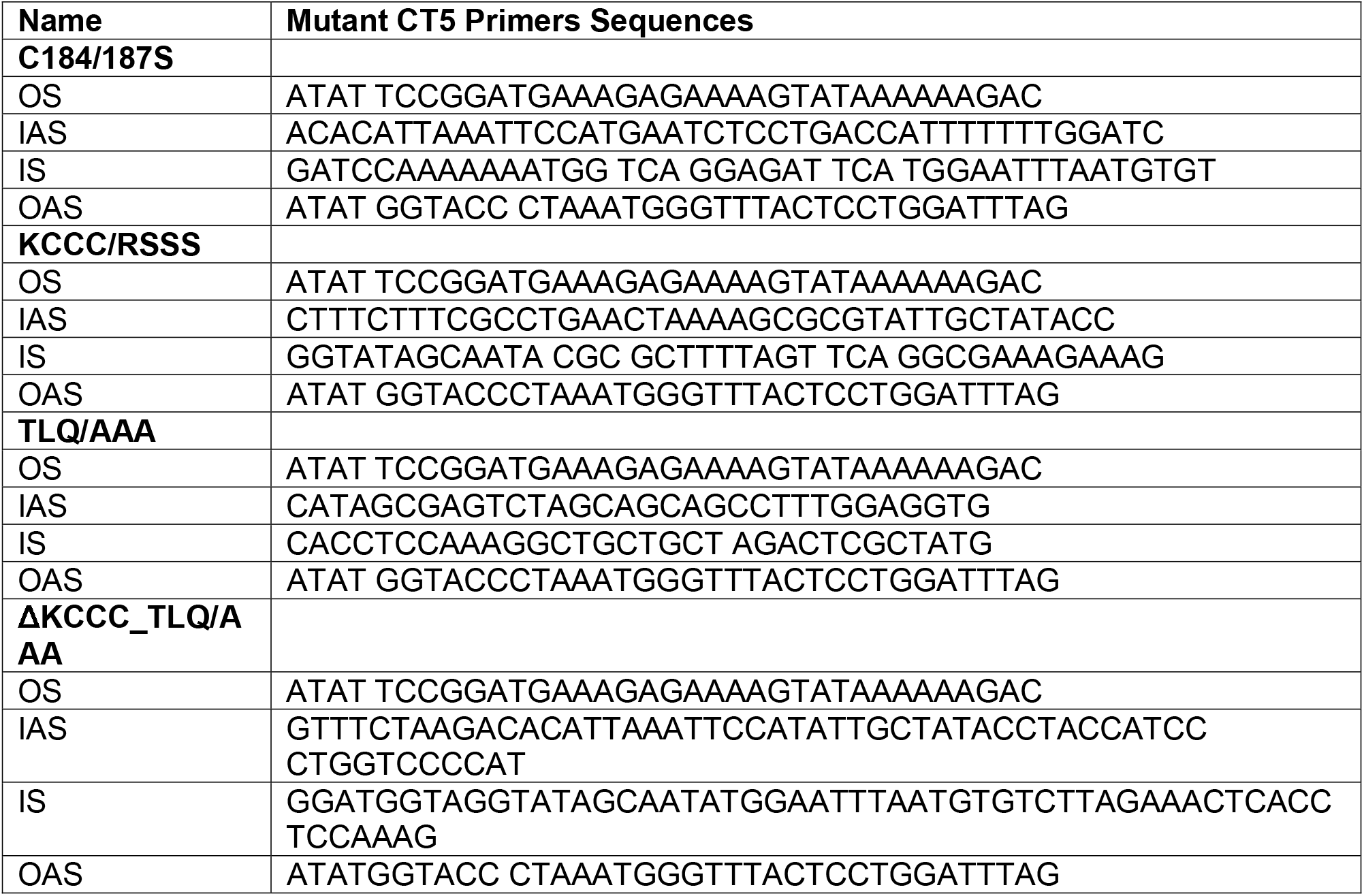
Overlap-Extension Primers to Generate Molecular Clones of Vif Mutant CT5.

#### Cloning, Expression and Purification of Recombinant FIV C36 Vif

Expression constructs containing FIV C36 Vif (residues 156–200 and 156–213) were generated by insertion into the EcoRI-Sbf I site of pMal c-5X (NEB). The resulting mannose-binding protein (MBP)-Vif fusion proteins were expressed and purified as previously described (59). Purified fusion proteins were concentrated in 20 mM HEPES pH 7.4, 150 mM NaCl, and 200 μM Tris (2-carboxyethyl) phosphine and used immediately for zinc-binding studies.

### ITC Analysis of Zinc Binding to Recombinant FIV C36 Vif

Isothermal titration calorimetry (ITC) experiments were performed at 25°C using a MicroCal iTC200 microcalorimeter (GE Healthcare) with a stirring rate of 1000 rpm. Complete details of the ITC assay have been described (59). Briefly, purified FIV Vif proteins were loaded into the ITC cell. A standardized stock solution of Zn(NO_3_)_2_ was loaded in the ITC syringe and injected step-wise into the cell. After the first injection of 0.4 μl Zn(NO_3_)_2_, 19 additional injections of 2 μL were made into the cell at 2 minute intervals to allow the system to return to equilibrium. Data were analyzed using MicroCal Origin software.

### Western Blotting

Following washing and trypsinization, cells lysates were obtained by suspending cell pellets in a volume of 10 μl per 100,000 cells with protease inhibitor (Complete-Mini, Roche) and an equivalent volume of 6X Laemmli buffer per sample. Lysates were boiled and clarified. Proteins were resolved on a sodium dodecyl sulfate–polyacrylamide electrophoresis (SDS-PAGE) gel and transferred to a polyvinylidene fluoride (PVDF) membrane (Thermo Scientific). Membranes were blocked in 10% milk and probed with primary antibodies. Membranes were incubated with appropriate secondary HRP-conjugated antibodies (dilutions 1: 5000–1:10000). Chemiluminescent substrates were used for visualization (Thermo Scientific 34080 or 34095).

### Western Antibodies

Rat anti-HA (Roche, high-affinity clone 3F10), rabbit anti-HA (Santa Cruz), mouse anti-HA (Sigma H3663), polyclonal rabbit anti-Myc (Santa Cruz), rabbit anti-tubulin (Santa Cruz Product 9104), and mouse anti-Flag (Sigma F3165) antibodies were used at a dilution between 1:500 and 1:1000.

### Proteasome Inhibition

Plasmid DNAs were transfected into 293T cells using calcium phosphate co-precipitation and medium was changed 12–16 hours later. Cells were treated with dimethyl sulfoxide (DMSO) or 10 □M MG132 (Sigma) 48 hours post-transfection, and cell lysates were harvested 22–24 hours later. Proteins were separated via SDS-PAGE as previously described.

### Co-Immunoprecipitation

Co-transfections of 293T cells were performed as above. Transfection media was replaced with DMEM 12–16 hours later. Then, 48–52 hours post-transfection, cells were washed with ice-cold PBS and lysed with ice-cold lysis buffer [150 mM NaCl, 50 mM TRIS HCl (pH 7.5), 1% Triton X, complete mini–protease inhibitor (Roche, Germany), and 200 mM phosphatase inhibitor (Sigma: Sodium Pervanadate)]. Sheep anti-mouse Immunoglobulin (Ig) magnetic dynabeads (sheep anti-mouse IgG, Invitrogen cat no.110.31; Oslo, Norway) were blocked with 0.01% bovine serum albumin (BSA) and 2 mM EDTA. Cell lysates were stored at −20°C. To eliminate non-specific binding, beads were incubated with cell lysates at 4°C for 1 hour. Beads from previous adsorption were discarded, and lysate were incubated with 1.0 μg mouse anti-HA antibody (Sigma H3663) to immunoprecipitate HA-tagged proteins. Pre-cleared lysates were incubated with 30 μl of pre-blocked dynabeads for 1 hour. Beads were washed three times with ice-cold co-immunoprecipitation lysis buffer. Immune complexes were eluted from beads with 30 μl of 2X β-mercaptoethanol Laemmli sample buffer and boiled at 95°C for 9 minutes. Proteins were separated by SDS-PAGE. Immunoblots were probed with a polyclonal rabbit antibody directed against the Myc epitope to assess the interaction between feline Cul5 Myc and FIV Vif HA-tagged (WT or mutant) proteins.

### Immunofluorescence

The 293T cells were seeded onto Nunc LabTEK II chamber slides (Thermofisher, Catalog Number 154461). Adhered 293T cells were transfected with Fugene (Roche). Then, 48 hours post-transfection, cells were fixed with 4% formaldehyde, washed with PBS, and perforated with ice-cold methanol. Cells were stained with primary antibodies against HA- (high-affinity rat anti-HA antibody; Roche) or Myc- (rabbit anti-Myc; Santa Cruz) tagged proteins and probed with secondary antibodies goat anti-rabbit Alexa Flour (AF) 488 or goat anti-rat AF 594 (Invitrogen) to analyze cellular distribution of proteins. Cells were stained with prolonged gold DAPI (Invitrogen) and mounted with a glass coverslip. All images were taken with a 510 laser-scanning confocal microscope (LSM 510).

### Single-Cycle FIV-Replication Assays

The 293T producer cells were co-transfected using the calcium phosphate method with the following plasmids: transfer (FIV human cytomegalovirus (CMV)–driven luciferase reporter) (60), vesicular stomatitis virus glycoprotein (VSV-G), FP93 packaging (express FIV: gag, pol, and rev, but lacks Vif and envelope proteins) (61), fA3s (either fA3Z3 or fA3Z2bZ3), and codon-optimized FIV C36 Vif-HA (WT or mutant; or empty vector parental control) plasmids. Then, 16–20 hours post-transfection, media was changed; supernatants were collected with a 0.45 μM filter 32 hours later. Freshly plated 293T cells were transduced with equal amounts of virus normalized to reverse transcriptase (RT). Media was changed 16–20 hours after infection. Seventy-two hours post-infection, supernatants were removed, cells were washed with PBS, and then lysed with 1% Tween. Quantification of luciferase activity was measured by the relative light units emitted from each sample. Error bars reflect standard deviation from the mean.

### Spreading Replication

The 293T producer cells were transfected with infectious molecular clones of WT or mutant 34TF10 Orf2 (CT5) using polyethylenimine (PEI) transfection reagent (Salk Institute). To boost first-round infectivity, vesicular stomatitis virus glycoprotein G (VSV-G) was co-expressed during production. Media was changed 6–8 hours later. Viruses were harvested, filtered (0.45 μM), and pelleted on a 20% sucrose cushion 48 hours post-transfection. Viral titers were determined using a focal infectivity assay (FIA). Freshly plated CrFk cells were infected, and 12–16 hours later, they were washed thoroughly to eliminate input virus. Every 72 hours, viral supernatant was harvested. Simultaneously, feline cells were divided, cultured, and passaged with fresh media. The RT activity of viral supernatants was determined as previously described (62).

### FIV Focal Infectivity Assay

The focal infectivity assay (FIA) assay has been previously described (63). In brief, CrFK cells were serially infected in 24-well plates. Sixteen hours post-infection, media was changed, and 40–42 hours post-infection, cells were washed and fixed with methanol. Cells were washed with 1X TNE (15.8 g Tris-HCl, 87 g NaCl, 7.44 g EDTA, 700 ml H_2_O pH to 7.5.), blocked (1X TNE, 10% FBS), and probed with PPR antisera (a generous gift from Peggy Barr) and goat anti-cat HRP secondary antibody (MP Biomedicals 55293). Cells were stained with aminoethyl carbazol (AEC Sigma A-6926) solution to visualize infected cells. Infected cells were counted with a hemocytometer to empirically determine the viral titer.

## Results

### Vif mutations affect fA3 protein degradation

Given that fA3Z2bZ3 is more active than fA3Z3 in preventing □Vif FIV replication (44–46), we used fA3Z2bZ3 in the majority of our experiments. We initially tested fA3Z2bZ3 with FIV Vif C36 harboring mutations in the BC box (TLQ/AAA, TLQ/TAQ, A205L, A205S, and A205T) and KCCC motif (K162R, C166S, C189S, C192S, KCCC/RSSS, and ΔKCCC) (Fig 1*A* and 1*B*). Wild-type (WT) and mutant FIV Vif proteins had similar intracellular patterns (Fig 1*C*) and expression levels (Fig 1*D*). In contrast, BC-box mutants TLQ/AAA, TLQ/TAQ, and A205L lost A3-degradative function (Fig 2A and B), while A205S and A205T maintained their function. Moreover, all KCCC mutants failed to direct fA3 degradation (Fig 2*C* and 2*D*). Treatment with the proteasome inhibitor MG132 partially rescued degradation of FIV Vif–mediated fA3Z2bZ3 and HIV-1 Vif–mediated hA3Z2g-Z1c (Fig 2*E* and 2*F*). Notably, Vif was poorly expressed when co-transfected with fA3Z2bZ3 and hA3Z2g-Z1c in DMSO-treated cells (Fig 2*E*). These findings suggest that Vif might be degraded when complexed with A3 proteins, consistent with rapid turnover of Vif and concomitant degradation when associated with A3 proteins (51, 64).

**Fig 2.**
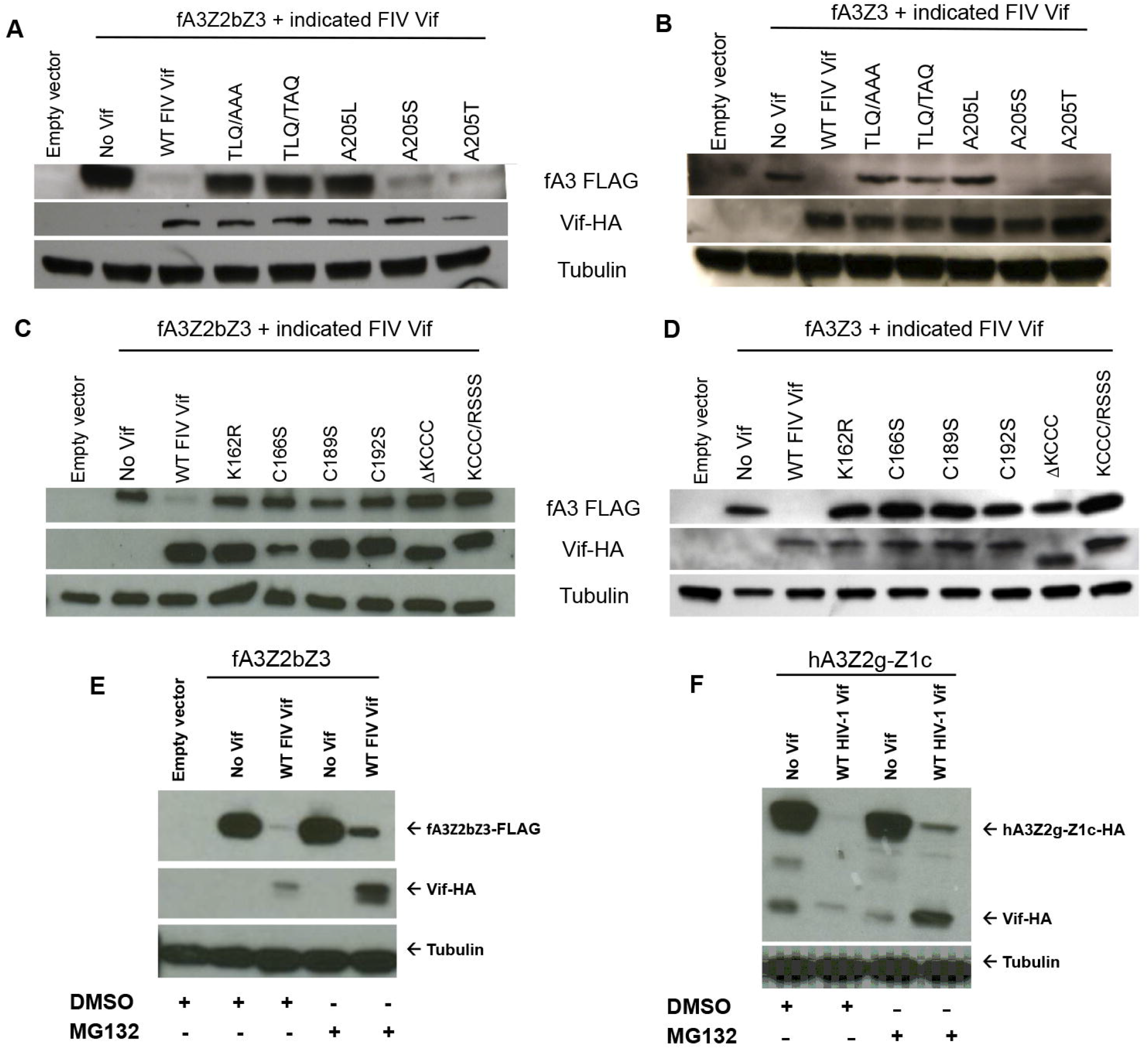
Effects of FIV Vif mutations on fA3 protein degradation. *A* and *B*, Specific FIV Vif BC-box mutants target fA3s for destruction. 293T cells were co-transfected with 0.5 μg untagged p1012, wild-type (WT) or mutant FIV Vif HA, and 1 μg of untagged pcDNA3.1(-) or FLAG-tagged fA3 (fA3Z3 or fA3Z2bZ3). Protein separation was analyzed by SDS-PAGE. Immunoblot membranes were probed with antibodies recognizing HA, FLAG, or tubulin proteins. *C* and *D*, FIV Vif with mutations in the BC box were unable to target fA3s for degradation. 293T cells were co-transfected with or without fA3s or untagged parental empty vectors with similar ratios of plasmid DNA as detailed in Fig 2*A* and 2*B*. A separate set of experiments showed FIV Vif mediates fA3Z2bZ3 proteasomal degradation. *E* and *F*, 293T cells were co-transfected with 0.5 μg of p1012 and 1 μg fA3Z2bZ3 or pcDNA3.1 (-) hA3Z2g-Z1c expression plasmids with 0.5 μg of WT FIV Vif or HIV-1 Vif with DMSO or MG132.

### Vif mutations influence viral infectivity

To determine whether loss of fA3 degradation correlates with increased restriction, we produced a virus containing a Δ*vif* FIV reporter both with and without fA3Z2bZ3 and the various Vif proteins. Consistent with their ability to target fA3Z2bZ3 for degradation (Fig 2*A*), A205S and A205T rescued FIV infectivity by 50% in the presence of fA3Z2bZ3 (Fig 3*A*). In contrast, TLQ/AAA, TLQ/TAQ, and A205L reduced FIV infectivity by 80% in the presence of fA3Z2bZ3 (Fig 3*A*). None of the KCCC mutants counteracted fA3Z2bZ3’s antiviral activity (Fig 3*B*). We confirmed the correlation between fA3 levels and infectivity with immunoblotting (Fig 3*A* and 3*B*). These data reveal novel residues within the BC box (TAQ, A205L, A205S, and A205T) and KCCC domain (K162R, C166S, C189S, C192S, KCCC/RSSS, and ΔKCCC) that are required for FIV Vif activity against fA3s and viral infectivity.

**Fig 3.**
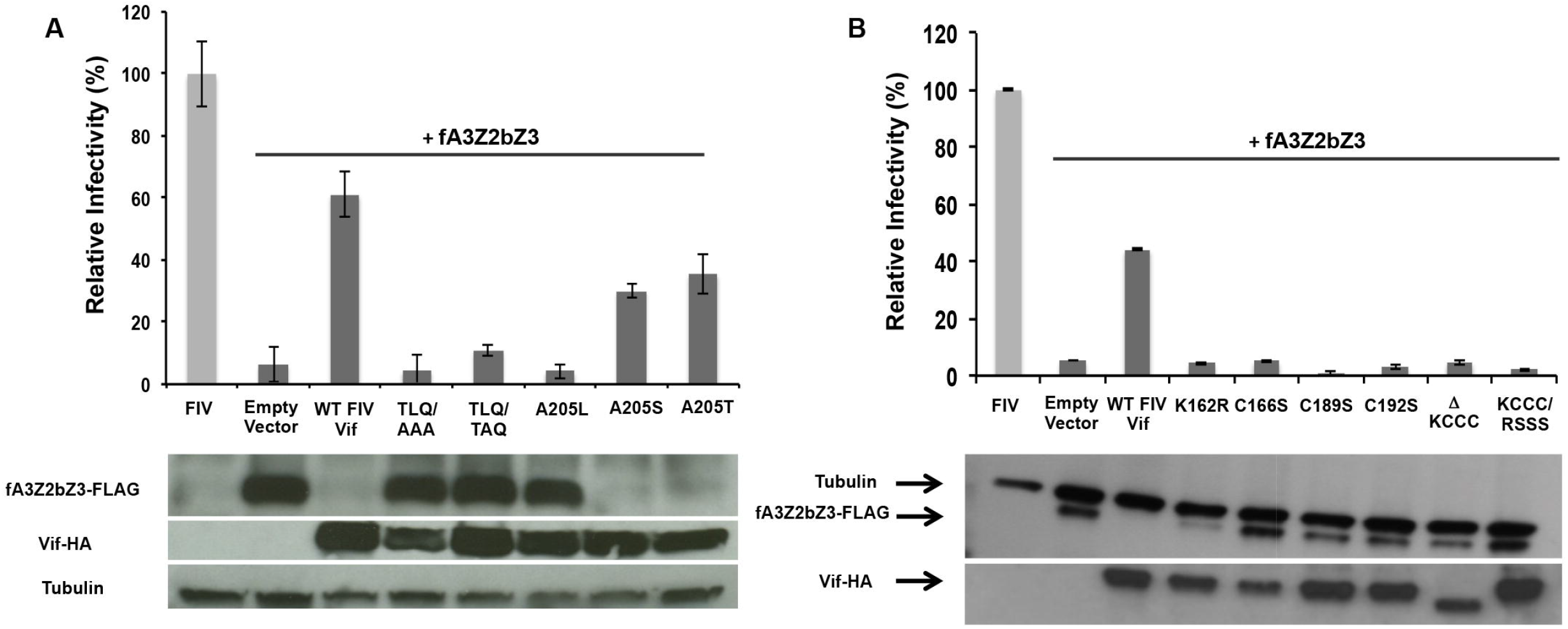
Effects of FIV Vif mutants on fA3CH restriction. *A* and *B*, Reporter viruses containing FIV ΔVif were produced in 293T cells transfected with 0.4 μg transfer vector, 0.2 μg pFP93 (FIV Gag-Pol and Rev) packaging vector, 0.2 μg pMD.G (VSV-G), with or without 0.3 μg of fA3Z2bZ3-FLAG and/or 0.5 μg Vif-HA [wild-type (WT) or mutant] plasmids, or the empty p1012 parental plasmid. The reverse transcription (RT) activity of each viral supernatant was determined, and equivalent amounts of RT-normalized virus were used to infect freshly plated 293T cells. Seventy-two hours after infection, lysates were analyzed for luciferase activity in triplicate. Standard deviation between triplicate samples was determined. FIV alone was set as 100%.

### Vif mutations alter spreading replication

We then introduced four Vif mutants (KCCC/RSSS, ΔKCCC+TLQ/AAA, TLQ/AAA, and C184+187S) into the full-length CT5 molecular clone to determine their effects on spreading viral replication. We chose these mutants because all of the KCCC mutants were defective for single-round infectivity. Stocks of mutant FIV virus were produced in 293T cells and tested in CrFk cells. All four mutants exhibited similar reverse transcriptase (RT) activities (Fig 4*A*), as 293T cells lack antiviral hA3 activity, and they did not negatively impact the infectious titer of the viruses (Fig 4*B*). Next, we infected CrFK cells with WT or mutant viruses at MOIs of 0.01 and 0.1. None of the four mutant viruses could replicate, while the WT virus replicated at 0.01 MOI, with large amounts of RT activity at 0.1 MOI (Fig 4*C* and 4*D*).

**Fig 4.**
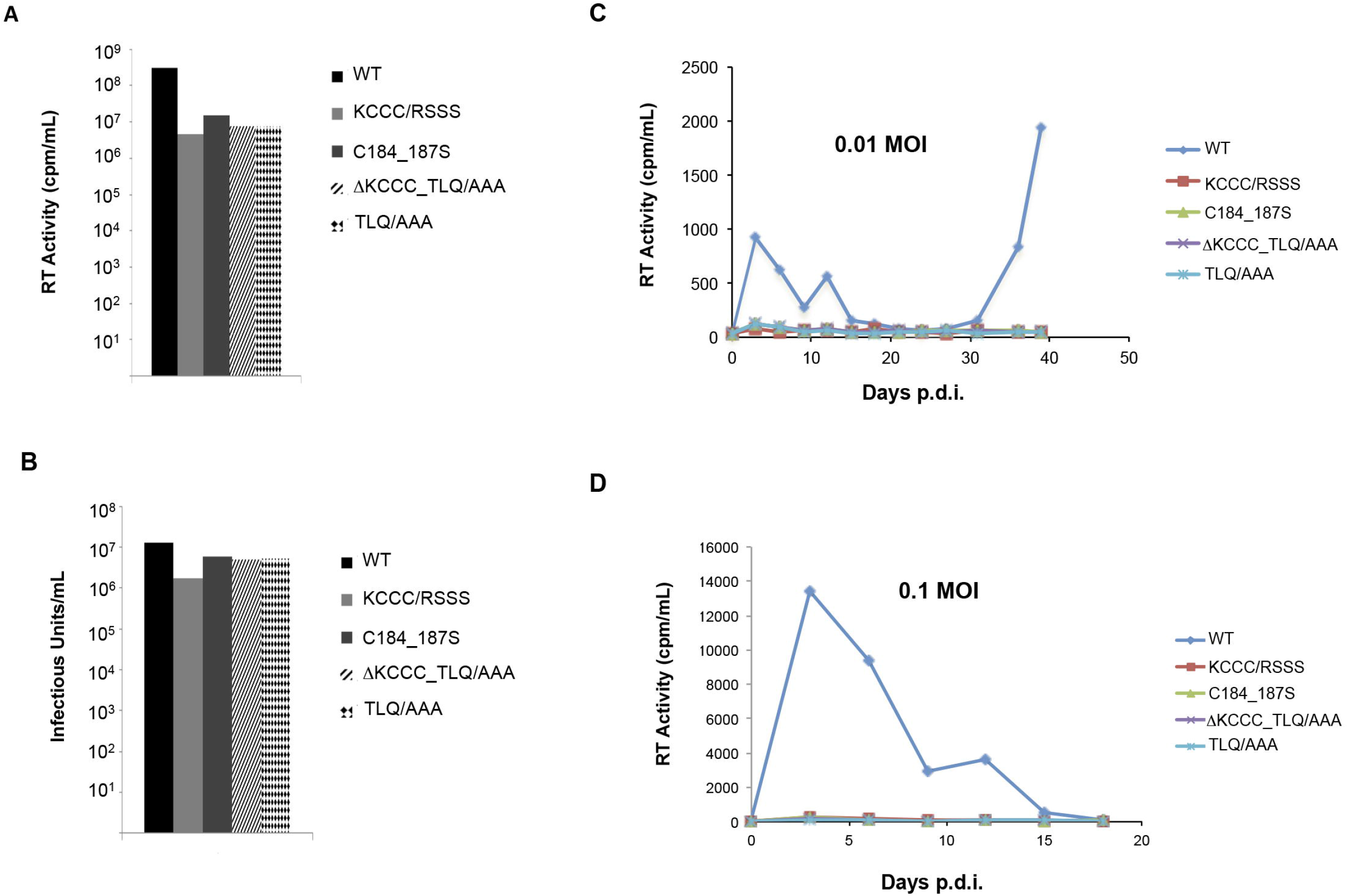
Effects of FIV Vif mutants block spreading replication of FIV. 293T producer cells were transiently transfected with VSVG-pseudotyped wild-type (WT) FIV or FIV Vif–mutant molecular clones (pCT5-based). *A* and *B*, Reverse transcription (RT) activity of viral supernatants and titers. *C* and D, Spreading replication: 4.5 × 10^4^ CrFK cells were infected at two different MOIs (0.01 and 0.1). Every 72 hours, viral supernatant was collected and stored at −80°C until RT activity was determined.

### Mutant Vif proteins co-localize and interact with feline Cul5

To date, the intracellular distribution of feline Cul5 (fCul5) has not been well characterized. Using confocal microscopy, we found fCul5 predominately in the cytoplasm, similar to Vif (Fig 5*A*). The KCCC mutants also display a slight intra-nuclear presence, in agreement with HIV-1 Vif, which has been shown to have a minimal Vif nuclear localization (65, 66). Co-immunoprecipitation studies assessed which FIV Vif domains were responsible for the interaction with Cul5. While the KCCC mutants interacted with fCul5 (Fig 5*C*, lanes 6–7), the TLQ/AAA mutant interacted poorly with fCul5 (Fig 5*C*, lane 8), as previously reported (51). Notably, when WT or mutant FIV Vif was co-expressed with fCul5 or hCul5, Cul5 expression significantly decreased (Fig 5*B*, lanes 5–8). As a result, less FIV Vif proteins were immunoprecipitated (Fig 5*C*, lanes 5–8), an effect not observed if hCul5 and HIV-1 Vif are co-expressed (Fig 5*B*, lane 4). Despite multiple attempts, we were unable to detect an interaction between feline Elongin B and C (fEloB and fEloC, respectively) in the presence of FIV Vif and fCul5. The latter may be due to reduced Cul5, which could have limited detection of the FIV Vif/E3 ligase constituents (e.g., fEloB and fEloC). Nonetheless, with these data, we verified the FIV Vif/fCul5 interaction reported by Wang and colleagues (51).

**Fig 5.**
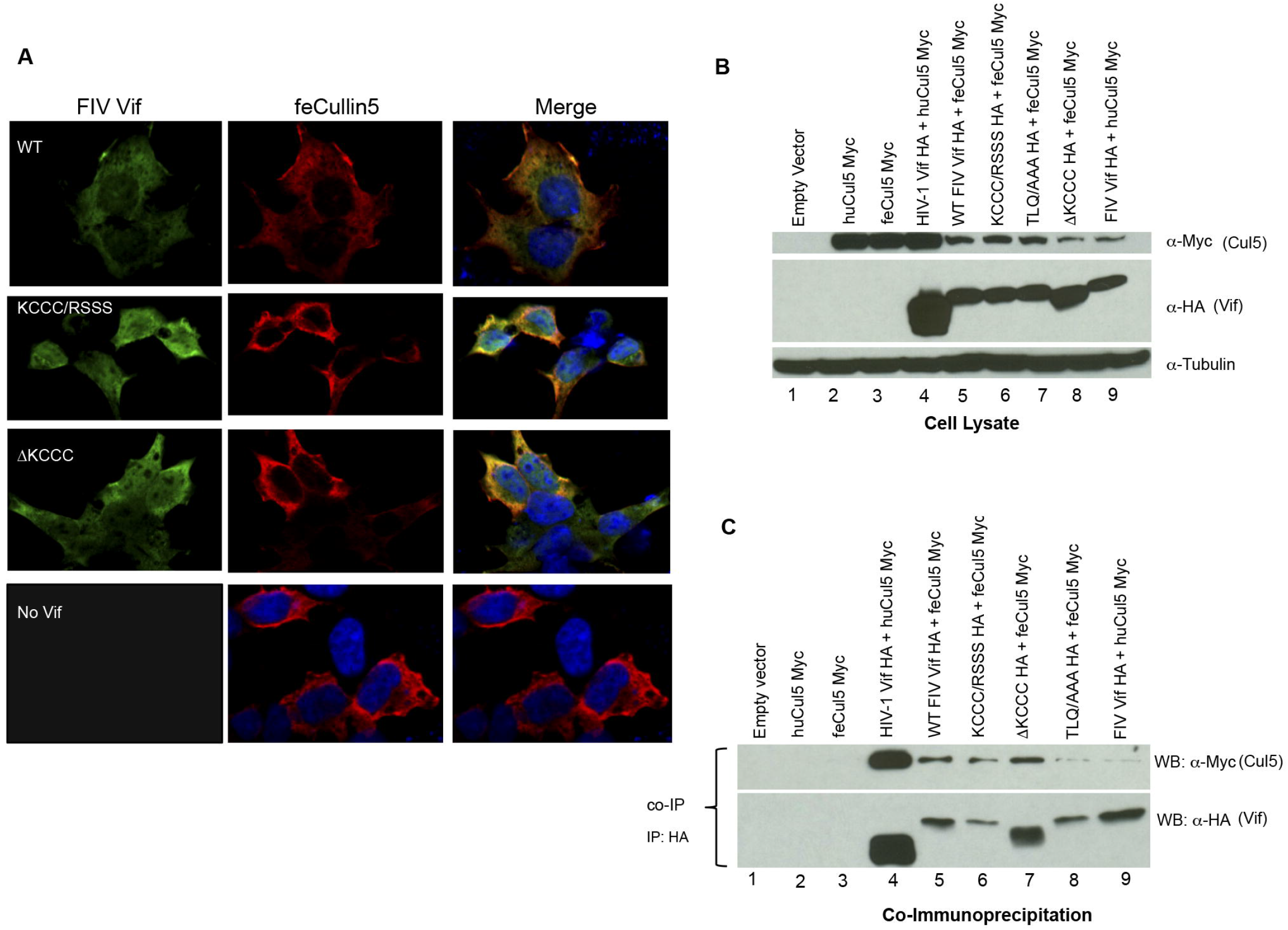
The FIV Vif KCCC domain does not mediate fCul5 interaction. *A*, 293T cells were co-transfected with 1 μg feline Cul5-Myc plasmid and 1 μg wild-type (WT) or mutant FIV Vif-HA plasmid using PEI. Then, 48 hours later, cells were immunolabeled and imaged using confocal microscopy. *B*, Cell lysates from 293T cells co-transfected with 3 μg of each expression plasmid were analyzed 48 hours post-transfection *C*, Protein-protein interactions. Lysates were immunoprecipitated with mouse anti-HA, and then incubated with sheep anti-mouse beads. Immune complexes were eluted and subjected to SDS-PAGE. Immunoblots were probed with a rabbit anti-Myc antibody to evaluate the interaction between fCul5-Myc and WT or mutant FIV Vif-HA proteins.

### Analysis of zinc binding to FIV Vif

A HCCH C-terminal motif in primate Vif proteins binds zinc and is required for Cul5 recognition. Feline Vifs have a similarly positioned KCCC motif. With isothermal titration calorimetry (ITC), we evaluated whether the KCCC domain of FIV Vif also binds zinc (59). FIV Vif proteins (Fig 6A) were expressed (Fig 6*B*) from the pMal-c5X vector and purified. All mannose binding protein (MBP)-Vif constructs showed no evidence of protein degradation (Fig 6*B*). Further, with size-exclusion chromatography, we separated and determined the protein quality of the purified MBP-Vif mutants. Notably, most of the N-terminal constructs did not bind to the amylose resin, suggesting that they may have formed aggregates that were unable to bind to the matrix (Fig 6*C*). Only two MBP-Vif mutants were monomeric (Fig 6*D* and 6*E*), as the rest formed aggregates (Fig 6*D*). Because protein aggregates can impair the biological function of a protein (67), the monomeric peaks that contained the intact KCCC-domain and BC-box motifs (156–213), as well as the domain that lacks the BC-box motif (156–200), were collected, concentrated, and analyzed further by ITC.

**Fig 6.**
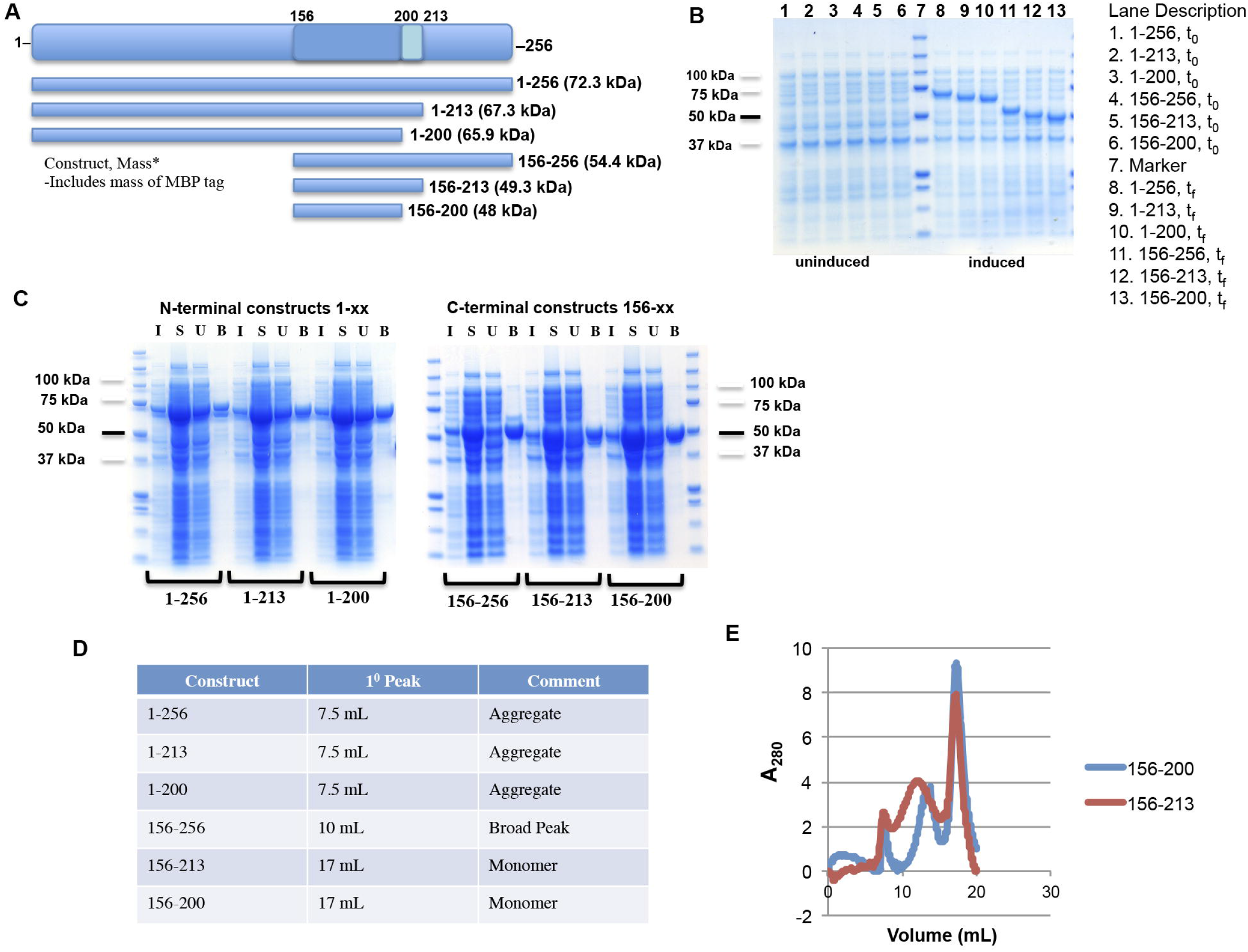
Expression, Purification, and Size Exclusion of FIV Vifs. *A*, Schematic map and numbering of purified and codon-optimized FIV_C36_ Vif. *B*, Expression of pre- and post-induction samples analyzed by SDS-PAGE. *C*, Verification of size exclusion chromatography SEC elutant and supernatant alongside pre- and post-induction samples were analyzed by SDS-PAGE. For each construct, the first lane is the induced (I) sample from Fig 6*B*, the second lane is the supernatant (S), the third lane is the unbound (U) flow-thru from the amylose column, and the fourth lane is the protein that bound (B) to and was eluted from the amylose column. *D*, Table of elution volumes and folding status of purified Vif MBP proteins. *E*, Monomeric resolution of purified Vif proteins (156–213 and 156–200) at 17 mL.

MBP-Vif proteins were concentrated in buffer containing excess reductant (tris-carboxyethyl phosphine) to maintain Cys residues in the thiol state. These thermogram data indicate that zinc failed to bind to either FIV Vif construct (Fig 7*A* and 7*B*), consistent with a recent study in which the zinc chelator PTEN failed to inhibit fA3 degradation by FIV Vif (51). These results suggest that FIV Vif must degrade fA3 proteins via a mechanism distinct from that of the zinc-binding primate Vif.

**Fig 7.**
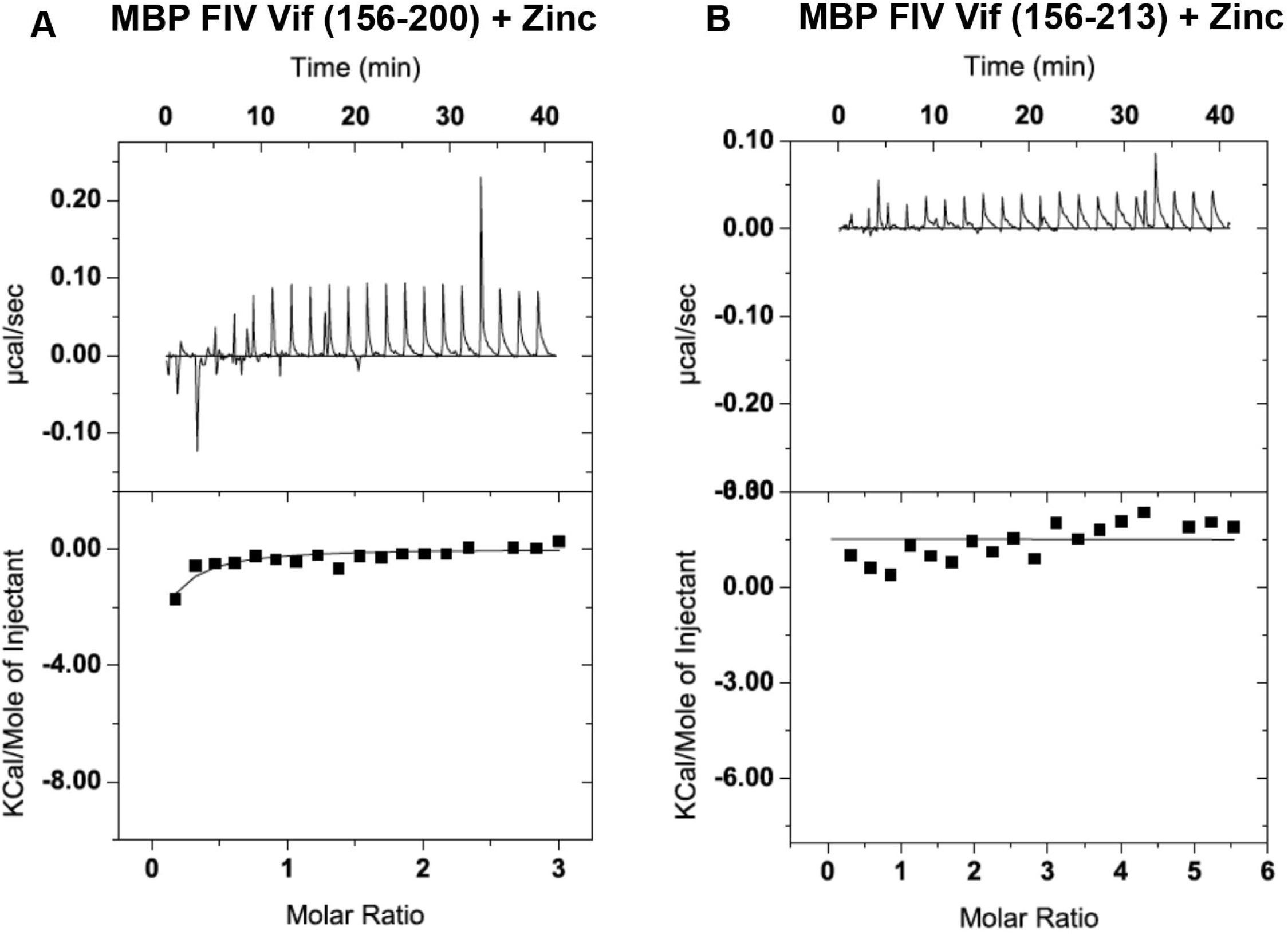
Isothermal titration calorimetry data for zinc titrations into FIV C36 Vif. *A* and *B*, Integrated heat data for MBP-Vifs (156–200 and 156–213, respectively).

## Discussion

With our data, we revealed novel residues within the FIV Vif BC box and KCCC domain that are required for infectivity and fA3 degradation. We also verified that the KCCC domain does not mediate the interaction with Cul5. In addition, we showed that FIV Vif does not bind zinc, and we are the first to demonstrate that FIV Vif (C36) targets human or feline Cul5 for degradation by an unknown mechanism. Interestingly, this phenomenon does not occur with HIV-1 Vif. These results collectively highlight a complex interplay between antiviral fA3 proteins, FIV Vif, and Cul5 that impacts FIV infectivity and replication.

Cullin proteins have an elongated structure that supports their conformational flexibility and facilitates binding to a broad range of substrate adaptors, including Vif. Vif usurps the host’s E3-ligase complex (which contains Cul5, Rbx, E2, Elongin B, and Elongin C) to steer A3 proteins to the proteasome. HIV-1 Vif contains a HCCH domain that coordinates zinc binding and mediates hCul5 binding (4, 8, 17, 19). However, the domain of FIV Vif that interacts with the fCul E3 ligase to target the two active anti-FIV fA3 proteins (fA3Z3 and fA3Z2bZ3) is not known (11). We hypothesized that residues in the KCCC motif, which has a similar location and composition as the HCCH motif in HIV-1 Vif, would bind to Cul5.

Prior to our studies, it was unknown whether introducing the BC box and KCCC mutants would disrupt the cytoplasmic distribution of Vif. Toward this end, we performed confocal microscopy, which showed that the KCCC and BC-box mutants were stably expressed (Fig 1*D*) and predominantly distributed in the cytoplasm (Fig 1*C*) with fA3Z2bZ3 (Fig 1*C*). The mutants co-localized with fA3Z2bZ3 in the absence of a proteasome inhibitor (Fig 1*C*). We believe that these FIV Vif/fA3Z2bZ3 complexes represent non-degraded populations that are incompletely targeted by the proteasome (51). We also tested whether previously unreported residues within the FIV Vif BC box would function similarly to HIV-1 Vif. Our results showed that two FIV Vif BC residues, TLQ/TAQ and A205L, were unable to target fA3 proteins for degradation (Fig 2*A* and 2*B*) as well as the reported TLQ/AAA mutant (48, 51). Interestingly, the A205S mutant targeted fA3s and partially rescued infectivity (Fig 2*A*, 2*B* and 3*A*). These results are consistent with the analogous A149S in HIV-1 Vif, which supports HIV-1 infectivity and hA3Z2g-Z1c elimination (15). In contrast, A205T restored FIV infectivity (Fig 3*A*), unlike the analogous A149T mutation in HIV-1 Vif (15). KCCC-domain mutants did not rescue Δvif FIV infectivity in the presence of fA3Z2bZ3 (Fig 3*B*), which correlates with their inability to mediate fA3Z2bZ3 degradation in virus-producing cells (Fig 3*B*). Collectively, these data support that the cytoplasmic localization of Vif proteins was not perturbed by mutations in the BC box and KCCC domain. Further, we demonstrated that both domains are necessary for targeting fA3s for degradation and single-round infectivity. We also provide data that conserving the alanine at residue 205 significantly contributes to the ability of FIV Vif to target fA3s for degradation. This finding suggests that the contact interface between Vif and fA3 proteins may not exactly mimic those within HIV-1 Vif and should be further studied.

An earlier report by Wang and colleagues showed that a triple alanine mutation (TLQ/AAA) and a KCCC/KCSS mutation in FIV Vif were able to block □Vif HIV-1 infectivity in human cells. Here, we aimed to determine the impact of FIV infectivity and spreading replication in human and feline cells, respectively. When we expressed a single-cycle, □vif reporter virus, FIV infectivity was incompletely restored, despite WT Vif being expressed in *trans*. Unfortunately, we found that studying FIV Vif complementation was technically challenging. In our hands, Δvif FIV infectivity in the presence of ectopically expressed Vif was only restored by 50–70%, despite multiple attempts to optimize the FIV packaging-VSVG-Vif-A3 plasmid ratios. Previous studies were able to complement □Vif FIV infectivity with expression of WT Vif in *trans;* however, different FIV Vifs that were derived from distinct isolates and packaging vectors were used (44, 46). Further, Wang and colleagues restored Δvif HIV-1 infectivity with WT FIV Vif_34TF10_ expressed in *trans* (51), while we used FIV_C36_ Vif. These experimental differences may explain the variation in restoring FIV infectivity between studies. Moreover, our FIV Vif mutants abolished spreading replication in CrFK cells, with a significant delay in FIV replication kinetics compared to WT Vif. Our data support previous work showing that Δvif FIV mutants are unable to restore FIV infectivity relative to WT controls in studies of spreading replication (68, 69). Further, we observed similar delays in spreading replication in CrFKs compared to peripheral blood mononuclear cells (PBMCs) derived from domestic cats and co-cultured with CrFK cells infected with Δvif FIV (69). CrFK cells express all fA3 proteins; hence, the delay in spreading WT replication may have been due to the antiviral activity of fA3 proteins in the absence of Vif (70). In support of this, all Vif mutant viruses were replication-incompetent, and the block in replication was sustained over 18 and 40 days at two different MOI (Fig 4*C* and 4*D*). Based on these data, the KCCC domain and BC box residues contribute to the essential function of Vif activity to enable FIV replication. Although, we do not formally show the mechanisms here, presumably our Vif mutants may be unable to overcome the restriction of fA3 proteins necessary to recruit an E3 ligase Cullin complex to steer these antiviral proteins to the proteasome.

Previously, localization of fCul5 was unknown. We found that fCul5 localizes to the cytoplasm, similar to hCul5 (Fig 5*A*), and that WT and mutant Vif proteins co-localize with Cul5 (Fig 5*A*). We determined that the BC-box mutant TLQ/AAA has a reduced interaction with fCul5 and examined whether the KCCC domain served as a binding motif for fCul5. By co-immunoprecipitation, we found that the KCCC mutants retained their ability to interact with fCul5 (Fig 5*C*), demonstrating that the KCCC motif is not required for the Vif-Cul5 interaction. While we cannot exclude that deleting the KCCC domain alters proper folding of FIV Vif, the KCCC/RSSS mutant produced similar results in that the KCCC/RSSS mutant retained its interaction with Cul5.

We verified that FIV Vif and fCul5 interact (Fig 5*B*, lane 5). Interestingly, we observed reduced fCul5 and hCul5 expression in the presence of WT and FIV Vif mutants, but not in the presence of HIV-1 Vif (Fig 5*B*, lanes 4–9). We consistently observed robust co-immunoprecipitation between HIV-1 Vif and hCul5 (Fig 5*C*). We suspect that an unidentified factor that is not expressed in fibroblasts may influence the stability of the FIV Vif/fCul5 complex.

Curiously, fCul5 and hCul5 are 99% homologous (51). However, Wang and colleagues did not observe reduced fCul5 expression in the presence of codon-optimized FIV_34TF10_ Vif in their interaction studies (51), as we showed with codon-optimized FIV_C36_ Vif in our co-immunoprecipitation studies (Fig 5*A*). Because the affinities of these protein-protein interactions are largely unknown, we question whether these two Vif variants differ in their ability to interact with fCul5. hA3s and HIV-1 Vif require different binding motifs to interact with fA3s and FIV Vif (71). Further, Troy and colleagues showed that the pathogenicity of FIV depends on the Vif isolate (70), which might correlate with the interaction, or lack thereof, between FIV_C36_ Vif and fCul5. Thus, our results demonstrate that a single residue within the BC box, TLQ/TAQ or A205L, of FIV Vif can induce fA3Z3 and fA3Z2bZ3 degradation, and that the KCCC motif does not mediate the interaction between fCul5 and FIV Vif.

Lastly, our biochemical ITC data showed that the KCCC domain of FIV Vif does not bind zinc (Fig 7*B*). We were unable to express a purified form of the full-length FIV Vif protein (Fig 6*D*), because purifying Vif is technically challenging (37, 72, 73). Nonetheless, our finding that FIV Vif does not bind to zinc agrees with studies of the PTEN zinc chelator carried out by Wang and colleagues (51). However, this result contrasts with the zinc binding we observed for HIV-1 Vif using ITC (59). Zinc binding to HIV-1 Vif changes the protein conformation and promotes interaction with Cul5. In the case of FIV Vif, such a conformational change is either not required or driven by other processes. Curiously, non-primate lentiviral BIV Vif requires zinc and has a novel zinc-binding domain, C-x1-C-x1-H-x19-C. Visna Vif, however, does not use zinc for its activity, but rather relies on a novel protein-protein interaction (74, 75).

While we were preparing this manuscript, Kane and colleagues showed that FIV_34TF10_ Vif interacts with both hCul2 and hCul5, albeit its binding to hCul2 requires the Vif N-terminal tag (75). Recruitment of Cul5 is mediated by the N-terminus of HIV-1 Vif (76), and stabilization between primate Vifs and Cul5 requires CBF-β, which targets hA3Z2g-Z1c for proteasomal degradation (27, 39, 72). Interestingly, FIV Vif does not require CBF-β to stabilize Vif, form the fE3-ligase complex, or mediate fA3-mediated degradation (39, 75), despite the identical homology between feline and human CBF-β (75). A non-primate Visna Vif has evolved to use a novel cellular co-factor, Cylophilin A, to promote the Visna Vif/Cul2 E3-ligase complex needed to degrade ovine A3 proteins, whereas BIV Vif does not require a co-factor to eliminate bovine A3 via the proteasome (75).

These studies illustrate the complex selection criteria that lentiviruses have undergone to allow species-specific Vif to adapt to host A3 proteins (71, 75, 77, 78). We hypothesize that the domestic cat may have flexible fCul2/fCul5 usage, unlike HIV-1 Vif, which predominantly interacts with hCul5. The reported study by Kane and colleagues focused on the FIV Vif–host interactome in human 293T cells (75). However, further studies in feline cells should determine whether Vif preferentially binds fCul2 or fCul5, examine if other feline Cullins interact with FIV Vif, and identify the elusive cellular co-factor that stabilizes FIV Vif. Based on our results, we believe that the lentiviral/host interactome hijacks novel cellular co-factors needed to eliminate the antiviral properties of A3 proteins, as is the case with FIV Vif, despite 100% homology between human and feline CBF-β (75). Data from these studies may serve as a platform for future experiments to provide mechanistic insight important for the development of novel antiviral HIV-1 therapies.

## Acknowledgments

We thank Dyana Saenz, and Mary Peretz for technical assistance and advice; Anne Meehan, James Morrison, and Hind Fadel for advice; Xiao Fang Yu (Johns Hopkins) and Jiawen Wang (Jilin University) for expression plasmids; and P. Barr (Western University of Health Sciences) for antisera. Supported by NIH AI100797.

